# Efficient overexpression and purification of SARS-CoV-2 Nucleocapsid proteins in *Escherichia coli*

**DOI:** 10.1101/2024.01.08.574531

**Authors:** Emma L Brudenell, Manoj B Pohare, Domen Zafred, Janine Phipps, Hailey R Hornsby, John Darby, Junxiao Dai, Ellen Liggett, Kathleen Cain, Perdita E. Barran, Thushan I de Silva, Jon R Sayers

## Abstract

The fundamental biology of Severe Acute Respiratory Syndrome coronavirus 2 (SARS-CoV-2) nucleocapsid protein (Ncap), its use in diagnostic assays and its potential application as a vaccine component have received considerable attention since the outbreak of the Covid19 pandemic in late 2019. Here we report the scalable expression and purification of soluble, immunologically active, SARS-CoV-2 Ncap in *Escherichia coli*. Codon-optimised synthetic genes encoding the original Ncap sequence and four common variants with an N-terminal 6His affinity tag (sequence MHHHHHHG) were cloned into an inducible expression vector carrying a regulated bacteriophage T5 synthetic promoter controlled by *lac* operator binding sites. The constructs were used to express Ncap proteins and protocols developed which allow efficient production of purified Ncap with yields of over 200 mg per litre of culture media. These proteins were deployed in ELISA assays to allow comparison of their responses to human sera. Our results suggest that there was no detectable difference between the 6His-tagged and untagged original Ncap proteins but there may be a slight loss of sensitivity of sera to other Ncap isolates.

## INTRODUCTION

The archetypal SARS-CoV-2 virus N gene encodes a nucleoprotein, also known as a nucleocapsid (Ncap) or N protein (N), consisting of 419 amino acids comprised of two domains (Uniprot entry P0DTC9)[1]. The N-terminal contains a predominance of beta-strand and coil structure with little helical content. In contrast to this, the C-terminal domain contains eight helical regions and a very short antiparallel 2-stranded beta-sheet feature (**Figure 1**) [2]. These domains are linked by a ∼ 70 amino acid unstructured linker region which appears to interact with non-structural protein 3 (NSP3) [3], a polypeptide consisting of 1945 amino acids which is involved in formation of viral replication-transcription complexes [4] as well as the membrane anchored viral M protein [5].

**Figure 1.**
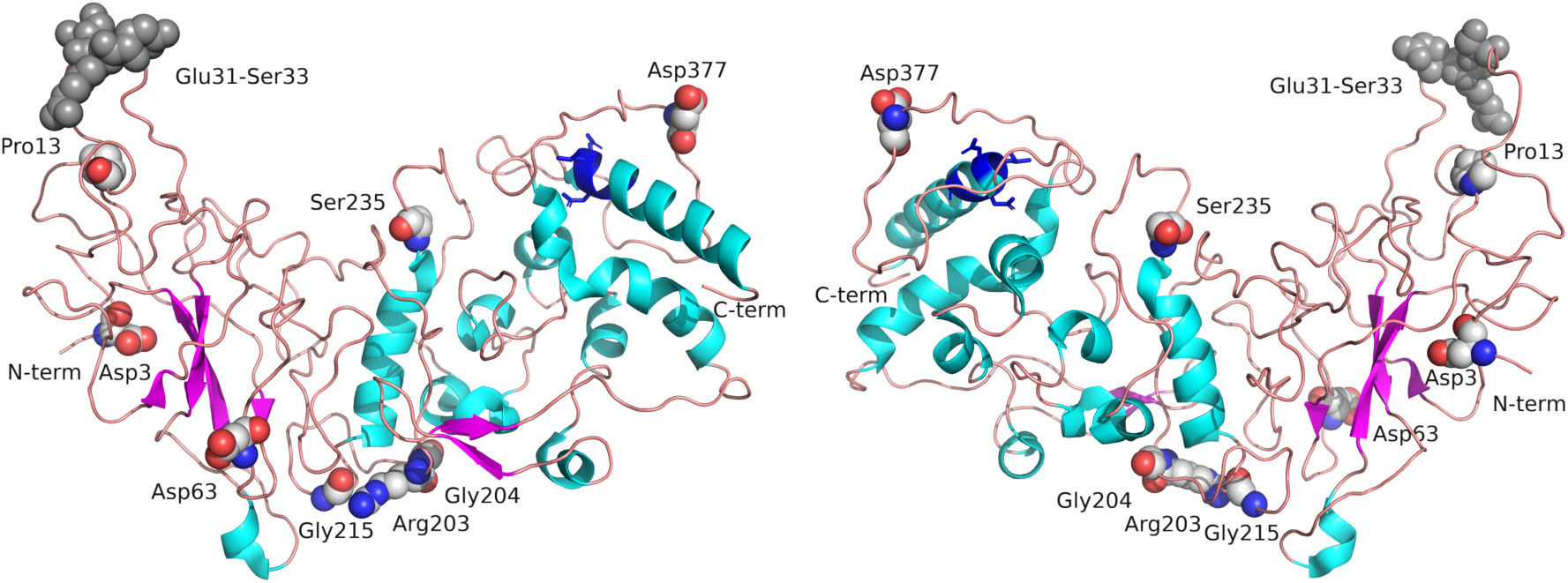
Structure of Nucleocapsid Protein. The structure of Ncap is shown rendered from 8FD5 in two views rotated 180° around a central y axis. Helical regions are shown in cyan and beta strands in magenta connected by loops possessing poorly-defined secondary structure. N and C terminal regions are labelled. Mutated residues in the proteins studied here are labelled and shown as spheres (carbon grey, oxygen red, nitrogen blue). The CASP6 cleavage site is shown in dark blue sticks (residues 399-402). Compared to the wild-type sequence, B.1.1 contains two amino acid variations, (Arg203Lys, and Gly204Arg), Alpha contains four (Asp3Leu, Arg203Lys, Gly204Arg and Ser235Phe), as does Delta (Asp63Gly, Arg203Met, Gly215Cys and Asp377Tyr) while Omicron has a deletion of Glu31-Ser33 (dark grey spheres) and two amino acid substitutions (Arg203Lys and Gly204Arg).

Both the N-terminal RNA-binding and C-terminal dimerization domains appear to interact with viral ribonucleic acid [5]. The protein is subject to post-translational modifications such as phosphorylation [6] and host-mediated proteolysis by the cysteine-aspartic protease caspase 6 [7]. This releases fragments that reduce the host’s inflammatory response by antagonising interferon gamma production [8].

Recombinant Ncap protein has also been used to facilitate development of both ELISA [9-12] and lateral flow diagnostics systems [12] [13] [14, 15]. The SARS-CoV-2 Ncap protein has been suggested as a possible vaccine candidate [16, 17] and it has been demonstrated that immunization with recombinant Ncap produced in *Escherichia coli* induced an antibody response in the lungs of rats [18]. Furthermore, next-generation vaccines are being developed that will express both Spike and Ncap proteins using adenovirus vectors [19] or using encapsulated mRNA approaches [20].

Since the first SARS-CoV-2 sequence was reported [1, 3], much attention has focused on mutations in the Spike protein and their impact on vaccine efficacy has been reviewed extensively [21, 22] [23]. However, mutations in the N gene encoding Ncap are also of interest as these may impact on pathogenicity [24] [25] and on diagnostic test efficacy. For example, the B.1.1 lineage [26] [27] arose in early 2020 which contained a double mutation in Ncap, Arg203Lys and Gly204Arg, [28] which spread throughout Europe and beyond. Further examples include Alpha (B.1.1.7) [29], Delta (B.1.617) [30] and Omicron (B.1.1.529) [31] amongst others [32] [33] [27]. Considerable ongoing sequence surveillance seems certain to identify further new variants [34] [35] [36] [37] [38]. Two studies report on whether Ncap protein sequence variation leads to reduced sensitivity for some rapid antigen tests (RATs). Hagag and co-workers found that the Arg203Met mutation, present in Delta, led to complete loss of detection {Hagag, 2022 #426] in the four RATs tests they examined, while a study examining eleven different commercially available tests found no cause to suspect that common variants circulating at the time would be less effectively detected [39].

Here we report the expression and purification and characterisation of recombinant Ncap proteins using codon-optimized synthetic genes in *E. coli*. Native (untagged) and Ncap adorned with an N-terminal tag consisting of six histidine residues were all produced in soluble form, purified and their responses to human sera compared. Tagged Ncap variants including B.1.1, Alpha, Delta and Omicron were also produced and examined.

## MATERIALS AND METHODS

### Bioinformatics and structure manipulation

Phyre2 [40] was used to model structures of Ncap based on a 5 Å CryoEM model 8FD5 [2]. The PyMOL Molecular Graphics System, version 2.5.2 Schrödinger, LLC was used to render 3D visualisations, to analyse mutations and for alignment of Phyre-2 generated structures and YASARA (YASARA Biosciences GmbH) was used to carry out energy minimisation of structures. The CABSFLEX server’s default parameters were used to compare protein flexibility in the variant sequences with the original by molecular dynamics simulations [41].

### Expression constructs

Synthetic genes or gene fragments were designed for optimal expression in *E. coli* (**Supplementary Table S1**). Genes were codon-optimised for the original SARS-CoV-2 nucleocapsid protein (Ncap, accession number YP_009724397) and its variants (B.1.1, QIQ08827; Alpha, QYU76755; Delta, UAL04655 and Omicron, UFO69287) in *Escherichia coli* using standard molecular biological methods [42]. Briefly, two synthetic gene fragments were purchased from Genewiz GmbH (Germany) and designated Ncap_*N-*terminal and Ncap*_C-*terminal. They contained a 25 bp overlap at their 3ʹ and 5ʹ ends respectively. They were mixed at an equimolar ratio and assembled into a complete coding sequence using SOE PCR amplification method [43] with either forward primer Ncap_for or Native_Ncap_for and reverse primer Ncap_rev (see **Supplementary Table S2**) to generate the N-terminal six-histidine tagged Ncap fragment or native construct lacking the tag. The resulting PCR products contain unique EcoRI and NdeI sites upstream of the coding regions and a downstream HindIII site as shown (**Table S1**). The PCR fragments were digested with restriction endonucleases HindIII and either NdeI or EcoRI, for native or 6His-tagged Ncap fragment, respectively and then cloned into expression vector pT5P digested with the corresponding enzymes.

This generated two clones designated pT5P_Ncap and pT5P_MHTNcap encoding the untagged and 6His-tagged proteins, respectively. Additionally, a variant encoding both Arg203Lys and Gly204Arg (B.1.1) with the N-terminal 6His-tag was produced by PCR using standard procedures. Briefly, the pT5P_MHTNcap plasmid was amplified with primers Ncap_for and Ncap_203/204_rev or Ncap_203/204_for and Ncap_rev (**Table S2**), to generate two fragments. These fragments were gel purified and mixed in equimolar ratio and PCR amplified with Ncap_for and Ncap_rev to generate 6His-tagged Ncap 203/204 fragment for restriction endonuclease-mediated cloning with *Eco*RI and *Hind*III into pT5P using standard methods generating the plasmid pT5P_MHTNcap_B11 which encoded B.1.1 Ncap sequence (Lys 203 and Arg 204). The Alpha and Delta, variants of Ncap, encoding the same min-His tag at their N-termini were custom synthesised (NBS Biologicals Ltd, UK) with codon optimisation flanked by EcoRI/NdeI and HindIII sites and supplied cloned in pUC57. The synthetic genes were inserted into expression vector pT5p using restriction enzymes EcoRI and HindIII generating plasmids pT5P_MHTAlphaNcap and pT5P_MHTDeltaNcap, respectively by standard methods [42]. The Omicron sequence was derived by inserting a synthetic gene fragment carrying the Omicron variation (custom synthesised by GeneWiz GmbH) between the NdeI and XbaI sites of pT5P_MHTNcap_B11, generating pT5P_MHTOmicronNcap. All insert sequences were confirmed by DNA sequencing by the University of Sheffield’s Core Genomics Facility or GeneWiz GmbH. Sequence of all the synthetic genes and fragments supplied and the proteins they encode are presented in **Table S1.**

### Protein expression

Production of recombinant protein from these plasmids was performed using standard methods. *E. coli* BL21 competent cells were transformed with the appropriate plasmid and grown on MDG agar plates [44] containing 100 µg/mL carbenicillin as follows. A freshly transformed single colony was used to inoculate 5 mL MDG media [44], supplemented with 1% vegetable-derived tryptone (Sigma 16922) and 100 μg/mL carbenicillin at 37°C, with shaking for 6 hours. The starter culture was used to inoculate fresh media containing 4% vegetable-derived tryptone, 2.5% yeast extract, 25 mM Na_2_HPO_4_, 25 mM KH_2_PO_4_, 50 mM NH_4_Cl, 5 mM Na_2_SO_4_, 2 mM MgSO_4_, 0.5%, glycerol (v/v), 0.05% glucose (w/v), and 200 μL of trace-metal solution per litre (Teknova, T1001). Typically, 500 mL cultures were grown in 2.5 L baffled flasks with a drop of antifoam (A6426, Sigma) at 37°C with vigorous shaking until they reached an absorbance of *A_600_* ∼3–4. Protein expression was induced by addition of isopropyl β-d-1-thiogalactopyranoside (IPTG, supplied by Melford UK) at a final concentration of 0.5 mM and the temperature lowered to 20°C for 24 hours to allow accumulation of the expressed protein. Alternatively, cells were grown in a small fermenter as follows: The fermenter was prepared using 10% vegetable tryptone and 5% yeast extract autoclaved in 2 L water. Separately autoclaved solutions of 100 mL 50xM (1.25 M Na_2_HPO_4_, 1.25 M KH_2_PO_4_, 2.5 M NH_4_Cl and Na_2_SO_4_), 10 mL 1 M MgSO_4_, 1 mL 1 M CaCl_2_ and 100 g 50% glycerol were added together with sterile filtered 50 mL 10% glucose, 5 mL carbenicillin (100 mg/mL) and 1 mL of trace elements solution (Teknova, T1001). The fermenter was topped up to 4.6 L with sterile water, warmed up, and 1 mL of antifoam (A6426, Sigma) was added just before the inoculation. A freshly transformed single colony was used to inoculate 5 mL MDG with carbenicillin (Studier) medium and grown at 37 °C to a density of A_600_ 0.9-1.0. It was then transferred to 400 mL MDG with carbenicillin and grown in a shake flask to a cell density of A_600_ 0.9-1.0 and used to inoculate the fermenter, which was stirred at ∼ 500 rpm and aerated with 5 L air per minute. The entire 400 mL inoculum was used to bring the volume to 5 L and cells were grown at 28 °C, doubling roughly every 60 minutes. Induction was carried out at a cell density ∼ A_600_ 3 by adding 0.5 mL of 1 M filter sterilised IPTG (final concentration 0.1 mM), and 10 mL of 10% lactose was added. Cells were harvested at a cell density equivalent to ∼A_600_=25, which corresponded to approximately 150 g of cell paste from a 5 L fermentation broth.

### Protein purification

Liquid chromatography was carried out on either an ÄKTA Prime or ÄKTA PURE system (Cytiva). Cell pellets (typically 10 g per batch) were resuspended in 50 mL in lysis buffer (25 mM Tris-HCl pH 8, 100 mM NaCl, 5% v/v glycerol, 1 mM 4-(2-aminoethyl) benzenesulfonyl fluoride hydrochloride) and lysed by the addition of 10 mg hen egg white lysozyme (Sigma, L6876) and sodium deoxycholate (Acros Organics) to a final concentration of 0.5 mg/mL then incubated overnight at 4°C. Subsequent procedures were carried out at room temperature. After sonication to reduce viscosity using short bursts (∼ 20-30 seconds) in an MSE Soniprep 150 Plus sonicator, the suspension was centrifuged at 30,000 x *g* for 30 minutes and the supernatant was recovered for immobilised metal affinity chromatography.

#### 6His-tagged Ncap proteins

The supernatant was adjusted to 5% (w/v) ammonium sulphate, mixed gently and centrifuged as above. The pellet was discarded. The supernatant was adjusted to 500 mM with solid NaCl and 17% w/v with ammonium sulphate to selectively precipitate the nucleocapsid protein at room temperature. The pellet was recovered by centrifugation as above and resuspended in 30 mL loading buffer (20 mM HEPES pH 7.8, 200 mM NaCl, 20 mM imidazole, 5% v/v glycerol) for purification by immobilised metal ion affinity chromatography (IMAC). Protein was loaded onto 2 X 5 mL HisTrap HP (Cytiva) columns connected in tandem and washed with wash buffer (20 mM HEPES pH 7.8, 2 M NaCl, 20 mM imidazole, 5% v/v glycerol) until the A_260nm_ /A_280nm_ ratio indicated bound nucleic acids were removed. Fractions (2 mL) were collected over a 10-column volume imidazole gradient (20 –500 mM, in 20 mM HEPES pH 7.8, 500 mM NaCl, 5% v/v glycerol). The purest fractions were diluted 10-fold into cation exchange loading buffer (20 mM HEPES pH 8, 10 mM NaCl, 1 mM EDTA, 1 mM DTT, 5% v/v glycerol) and loaded on to a 20 mL HiPrep SP FF 16/10 column (Cytiva). Protein was eluted over a linear gradient (10 – 1000 mM NaCl) and 2 mL fractions were collected. Samples were analysed by SDS PAGE on 10% polyacrylamide gels. The purest fractions were collected and buffer exchanged with storage buffer (20 mM HEPES pH 8, 120 mM NaCl, 1 mM EDTA, 1 mM DTT, 10% v/v glycerol) by ultrafiltration using Amicon Ultra–15 10 kDa cut-off centrifugal filters and flash frozen in liquid nitrogen for long-term storage at -80°C.

#### Untagged Ncap protein

Cells were lysed as above except that the buffer also contained 1 mM EDTA. After centrifugation the cell lysate was subjected to sonication to reduce viscosity and adjusted to 5% (w/v) in ammonium sulphate. Sufficient 5% polyethyleneimine-HCl (PEI, pH 8) was added to precipitate nucleic acids and mixed gently on a roller for 30 minutes prior to centrifugation at 30,000 x *g* for 30 minutes to remove the precipitated RNA/DNA/PEI pellet. Ncap protein was selectively precipitated by gradual addition of solid ammonium sulphate with gentle stirring to a final concentration of 17% (w/v) in presence of 500 mM NaCl. The pellet, consisting largely of precipitated Ncap was recovered by centrifugation as above and resuspended in 200 mL of cation-exchange loading buffer (20 mM HEPES pH 8, 10 mM NaCl, 1 mM EDTA, 1 mM DTT, 5% v/v glycerol) and loaded on to a 20 mL HiPrep SP FF 16/10 (Cytiva) column. Protein was eluted over a 20-column volume linear gradient (10 – 1000 mM NaCl) and 2 mL fractions were collected. Samples were analysed by SDS PAGE on 10% polyacrylamide gels. The purest fractions were concentrated and loaded on to a HiLoad Superdex 200 16/600 120 mL (Cytiva) column for size exclusion chromatography and eluted in 25 mM Tris pH 8, 120 mM NaCl, 1 mM EDTA, 5% glycerol (v/v). The purest fractions were collected and stored at -80°C by flash freezing in liquid nitrogen.

### Mass Spectrometry analysis

Analytical services were provided by the Faculty of Science biOMICS Facility, and Dept of Chemistry, University of Sheffield, and the Michael Barber Centre for Collaborative Mass Spectrometry, University of Manchester UK. All protein samples were buffer exchanged into 50 or 200 mM ammonium acetate (Fisher Scientific 10365260) as indicated using Zeba Micro Spin columns (#89877,75µL) prior to mass spectrometry analysis. Intact mass analyses were performed on a Waters Vion IMS Qtof connected to an Acquity I-Class liquid chromatography (LC) system. Protein samples were separated using an Acquity UPLC BEH C4 column (p/n 186004496), maintained at 80°C. The system was controlled using UNIFI software in positive ion mode with a capillary voltage of 2.75 kV and source temperature of 150°C. The LC gradient was developed at a flow rate of 0.2ml/min over 10 mins as follows: 0 min: 5% B, 1 min, 50% B 3.5min: 95% B 7.5min, 5% B, 10min (mobile phases A: water (MQ)/0.1% formic acid and mobile phase B (100% acetonitrile/0.1% formic acid). Data analysis was performed using UNIFI. Native mass spectrometry was carried out using direct-infusion nano-electrospray ionisation from in-house pulled borosilicate capillary tips on a Waters Synapt G2-S in positive ion mode, capillary voltage of 1.1-1.4 kV, desolvation temperature of 80°C and cone voltage of 10V. Data analysis was performed using MassLynx and Origami software.

#### Serology

The IgG-specific responses of pooled anonymised serum samples from hospitalised patients with confirmed COVID-19 (obtained with approval from the Sheffield Teaching Hospitals’ Research and Development office (Sheffield, UK)) to Ncap proteins were assessed by ELISA assays exactly as reported previously [45]. Ncap proteins were immobilised in microtitre plates (Immulon 4HBX; Thermo Scientific, 6405) at 4 °C overnight at 2 μg/mL (50 µL per well) in PBS (pH 7.4). Plates were washed with 0.05% PBS-Tween, then blocked for 1 hour at room temperature (RT) with 200 μL/well 0.5% casein buffer. The IgG response curves were generated by serially diluting in 1·75× steps from an initial dilution of pooled serum at 1:200, 100 µL loaded per well, and incubated for 2 hours at RT, followed by washing and addition of 100 µL/well of goat anti-human IgG-HRP conjugate (Invitrogen, 62-8420) at 1:500 dilution, and incubation for 1 hour at RT. Wells were then washed and 100 µL/well TMB substrate (KPL, 5120-0074) was added and left to develop for 10 minutes. Stop solution (KPL, 5150-0021) was added at 100 µL/well, and the absorbance read at 450 nm. The WHO International Standard for anti-SARS-CoV-2 immunoglobulin (NIBSC, 20/136) was used for calibration of this convalescent serum pool and readings are thus expressed in WHO binding antibody units per millilitre (BAU/mL).

## RESULTS & DISCUSSION

### Potential impact of mutation on Ncap function

We modelled the impact of mutations on local protein structure (**Figure 2** and **S1**). Unsurprisingly, deletion of residues 31-33 in the Omicron variant results in the largest apparent structural changes (**Figure 2**) but the Pro13-Leu substitution appeared to cause little local structural perturbation to the modelled, energy minimised structures. Other substitutions also seem likely to have an obvious impact. For example, the Asp3Leu and Ser235Phe substitutions in Alpha Ncap result in loss of two charge-charge (ion pair) interactions with arginine residues 88 and 92 in the former as well as loss of a hydrogen bond between the serine hydroxyl group and backbone carbonyl of amino acid 189 in the latter (**Figure S1A**). Similarly, the Arg203Lys/Gly204Arg double substitution present in B.1.1, Alpha and Omicron, potentially result in additional hydrogen bonds formed between the side chains of Arg204 (with amide oxygen of residue Ser202) and Lys203 (the backbone of Asp215) as shown (**Figure S1B**). In contrast, amino acid substitutions Asp63Gly and Arg 203Met in Delta Ncap both result in loss of a charged partner in ion pair interactions (with Lys65 and Asp216, respectively) compared with WT Ncap as (**Figure S1C**).

**Figure 2.**
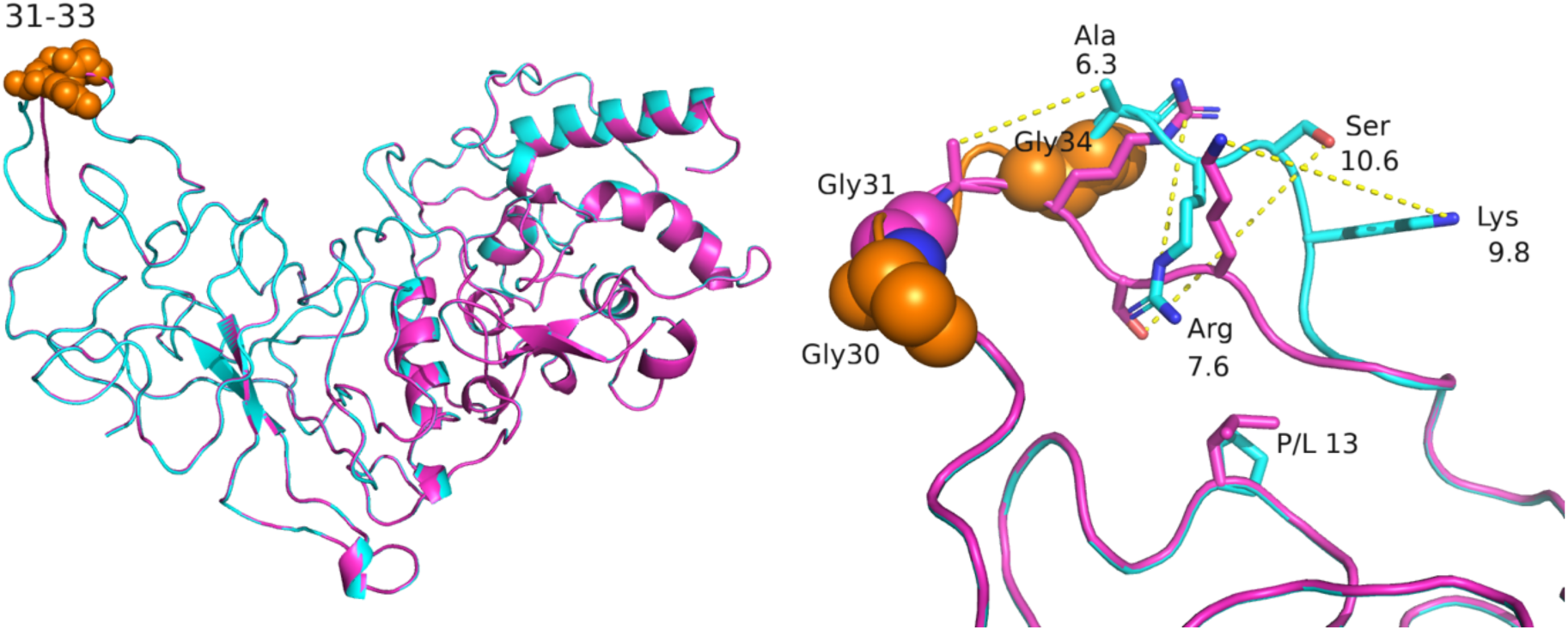
Impact of Omicron variant mutations on Ncap structure. Wild type Ncap structure (cyan cartoon, based on PDB entry 8FD5) was aligned with the predicted structure of the Omicron variant (magenta) as shown in the left panel. The two structures overlay well except in the region of the Omicron deletion. In Omicron, the deletion of residues 31–33 results in two adjacent glycine residues (residue 30, orange spheres and 31, previously 34 in the WT, magenta spheres). The side chains of residues Ala-Arg-Ser-Lys (35-38 in the WT sequence) show the largest structural changes (dotted yellow line show displacement in Å). Mutation of proline 13 to leucine (P/L 13) is predicted to have minimal structural impact.

In addition to the full-length CryoEM structure [2], several X-Ray structures have been reported for isolated N or C terminal domains [46] [47] [48] [49] [50], and one NMR structure of the “linker region”, residues 191-262 [3] which shows interactions with the viral protein NSP3.

Several of the mutations in Ncap map to this linker region so we examined how they might affect these interactions with NSP3. This latter structure includes residues Ser235, Arg203, Gly204, Gly201 and Ser235 but there would appear to few direct NSP3 interactions (**Figure 3A**) as they mostly are located at a considerable distance from the NSP3 protein as shown in the NMR ensemble (Figure 3A). Ser235 is a possible exception to this (**Figure 3B**). In one conformation of the complex, this residue approaches NSP3 to within 5 Å of Lys38. In Alpha Ncap, this residue is mutated to a phenylalanine. We modelled this change into the complex which was then energy minimised revealing a possible π-cation interaction between the phenylalanine and the epsilon-amino group of the lysine residue. Depending on the environment, π-cation interactions can be stronger than salt bridges, providing some potential stabilization of the complex [51] in this mutant.

**Figure 3.**
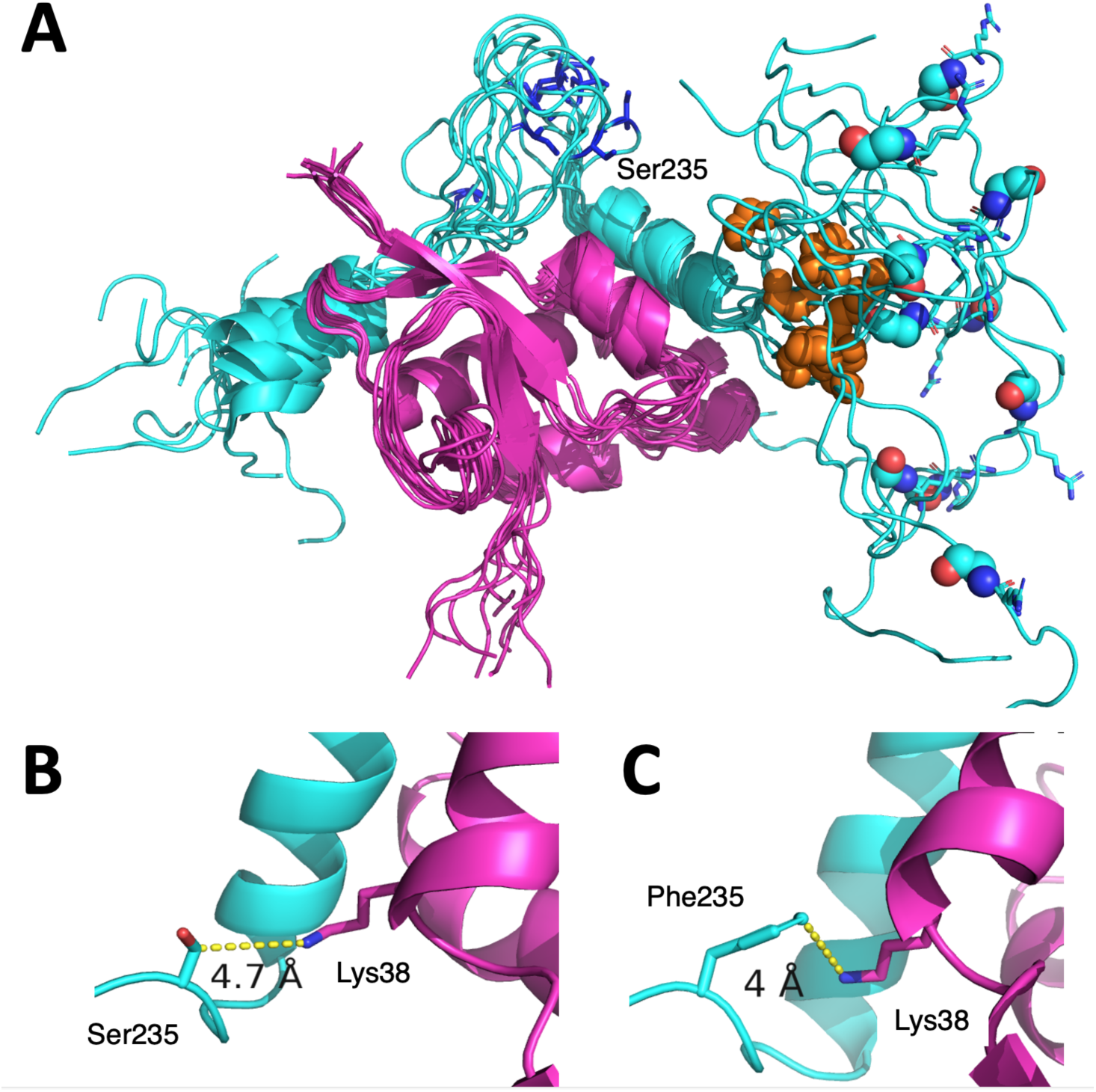
Interactions between the Ncap linker region and NSP3. **(A)** The ensemble NMR structure of Ncap (cyan cartoon) in complex with NSP3 (PDB code 7PKU). Glycines 204 and 215 are shown as spheres, in cyan and orange respectively. Ser235 is labelled and shown as blue sticks and Arg203 shown as cyan sticks. **(B)** Ser235 approaches closest of all residues in the ensemble, coming to within 5 Å of NSP in one confirmation as shown. **(C)** A potential π-cation interaction between the Phe235 mutation present in the Ncap Alpha variant was modelled.

Mutations in Ncap could alter its ability to bind nucleic acid. However, predicting the impact of amino acid substitutions on such interactions is made difficult as only limited information on the structure of Ncap bound to RNA is available. A structure of residues within the isolated N-terminal domain, (residues 48 – 171 of pdb code 7XWZ) bound to RNA does not indicate any direct interactions between the altered amino acids discussed above and the nucleic acid bound as they are not present in the structure determined by CryoEM [2]. However, as both N and C-terminal domains are implicated in RNA binding, which involves dimerization/oligomerization, it is possible that mutations in Ncap could alter interactions with their substrates [52].

We carried out preliminary molecular dynamics simulations to gain insight into potential impact the amino acid substitutions might have on Ncap flexibility (Figures S2&3). Analysis of an ensemble of structures obtained by molecular dynamics simulations [41] for each of the variant Ncaps revealed no large changes to the structural stability of the proteins. However, some differences between the original Ncap and variants were detected. The sum of the atomic root-mean-square fluctuation across the MD trajectory for Ncap and each variant were 701 Å for Ncap, 725 Å for the B.1.1, 650 Å for Alpha, 685 Å for Delta and 621 Å for Omicron. These variations are shown in **Figures S2&3** along with differences between predicted MD trajectories for each residue in the variants compared with the original sequence.

### Protein production

Ncap variants were cloned and expressed using the pTT5 expression vector developed in our laboratory which was derived from plasmid pTTQ18 [53]. It has a strong but inducible bacteriophage T5 promoter [54] under the control of the *lacI^q^* repressor which is also encoded on the plasmid. As the T5 promoters are recognised by *E. coli* RNAP polymerase, this system can be used in most commonly available host strains. We used BL21 [55] for the work described here in combination with media based on that described by Studier for the expression of the Ncap proteins [44]. All nucleocapsid proteins were expressed in soluble form and the purification procedure outlined above produced proteins which were >95% pure as exemplified for the Omicron variant (**Figure 4**, others shown in **Figure S4**). This contrasted to previous reports in which recombinant SARS-CoV-2 nucleocapsid protein was obtained as insoluble aggregates in *E. coli* requiring purification under denaturing conditions followed by refolding [56] [57] [58] or required fusion with maltose-binding protein to aid production of soluble material [59] followed by proteolysis of the fusion partner. Ncap and variants were obtained at yields of 7–10 mg or more of purified protein per g of cell mass from fermentations yielding 20-30 g of cells per litre of culture grown in modified Studier medium [44].

**Figure 4:**
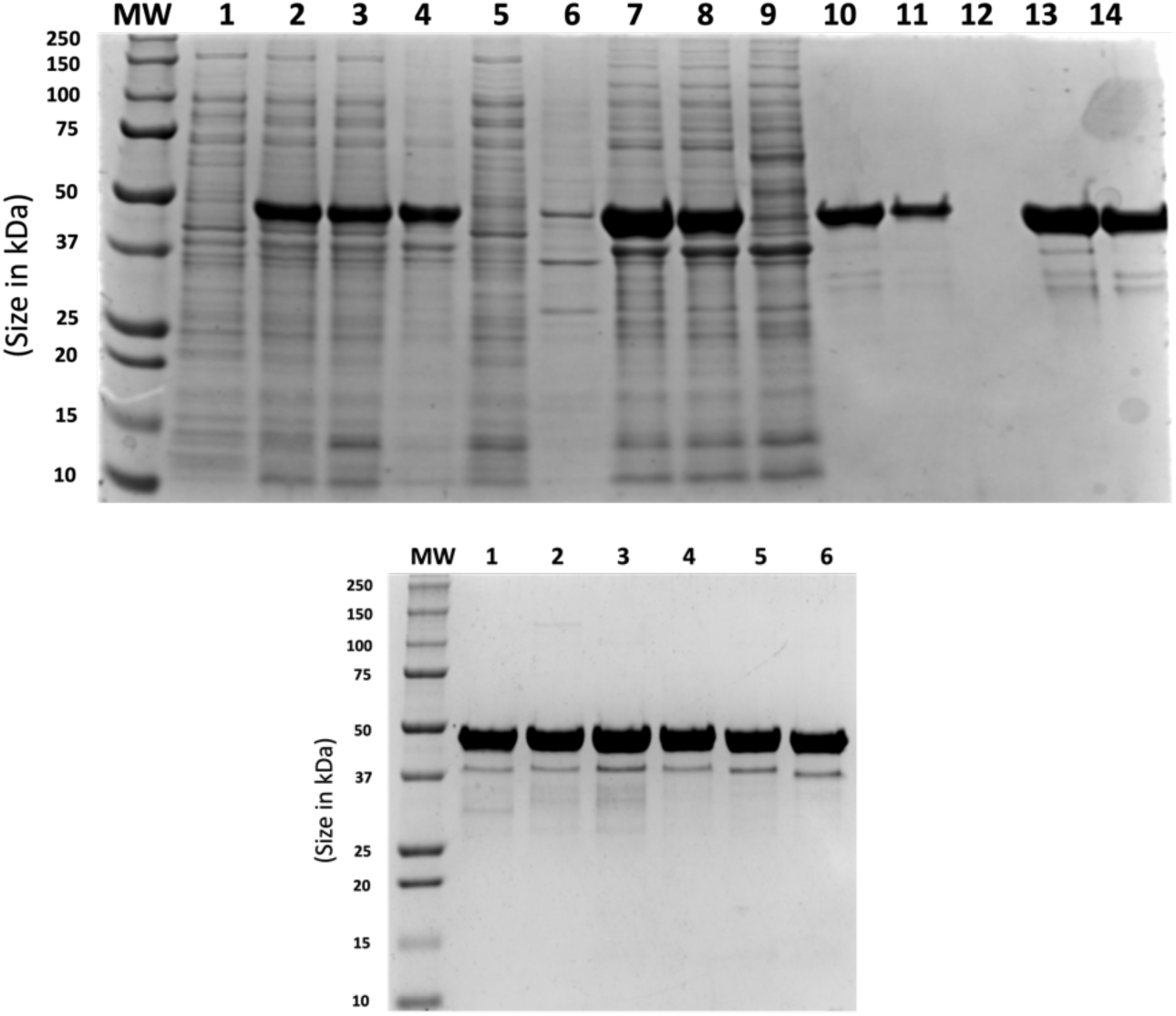
Expression and purification of 6His-tagged SARS-CoV-2 Ncap Omicron. SDS-PAGE (10%) analysis of samples from example purification. Molecular weight markers (MW, BioRad (Cat. 161-0362)) are shown for each panel. Left panel shows total SDS-cell lysates of uninduced and induced *E. coli* BL21 carrying plasmid pT5P_MHTOmicronNcap (Lanes 1 and 2 respectively). The soluble fraction from induced cells after lysis, sonication and centrifugation (Lane 3). Pellet from 5%-17% ammonium sulphate precipitation (Lane 4) and corresponding supernatant (Lane 5). The ammonium sulphate-protein pellet was resuspended in HisTrap loading buffer, centrifuged to separate into insoluble material (Lane 6) and the soluble fraction (Lane 7) and loaded on a HisTrap column (Lane 8). Lane 9 shows proteins passing through the HisTrap column. Pooled fractions eluted from the HisTrap column (Lane 10) were diluted with cation exchange loading buffer and loaded on SP Sepharose column (Lane 11). Flow through from SP column (Lane 12). Peak fractions from linear gradient elution (Lane 13 and 14). Right panel shows samples of purified 6His-tagged; Ncap Omicron (Lane 1); Ncap Delta (Lane 2); Ncap Alpha (Lane 3); Ncap 203/204 (Lane 4); Ncap (Lane 5) as well as untagged Ncap (Lane 6).

A differential ammonium sulphate precipitation (5% -17% w/v) removed significant quantities of *E. coli* host cell proteins before further liquid chromatography steps. Further purification was carried out by IMAC followed, in the case of the 6His-tagged proteins and cation exchange chromatography by elution with imidazole or salt gradients, respectively.

All proteins were shown to be free of significant nucleic acid contamination as shown by UV spectroscopy (**Figure S5**). We noticed that Ncap proteins had a tendency to co-purify with nucleic acids as monitored by the samples’ A_260nm_/A_280nm_ ratios. It was particularly important to wash the 6His-NCAP proteins bound to the IMAC column with sufficient high-salt buffer to remove the co-purifying nucleic acids, requiring up to 20 column volumes depending on the sample, or in the case of the native protein, using PEI precipitation for the same purpose. Mass spectrometry (electrospray) showed that each protein displayed the expected mass within experimental error (**Supplementary Table S3**).

### Identity of minor contaminants in recombinant Ncap

As we required our Ncap proteins for immunological assays, we were keen to understand the level of purity obtained and gain insight into the nature of the inevitable contaminating *E. coli* proteins which would be present, albeit at low levels. Thus, a more detailed analysis of our 6His-tagged and untagged Ncap proteins was carried out to identify the contaminating bands that became apparent when purified samples were over-loaded on SDS-PAGE gels. **Figure S6** shows SDS-PAGE gels with overloaded purified samples. Minor bands with both higher and lower molecular weights than the main expected product were observed. However, when these bands were excised from the overloaded gels and subjected to proteomic analysis via tryptic digest with quantification estimated using intensity based absolute quantification (iBAQ) [60] all the contaminant bands appeared to have the recombinant nucleocapsid as their main constituent (>90% for all minor bands analysed). In both the untagged and tagged Ncap, the main band on the gel was comprised of over 97% recombinant Ncap as estimated by iBAQ. Human keratins, proteomic workflow contaminants, were also observed but were excluded from the analysis. The observation that the minor high-MW contaminants observed on SDS-PAGE at high loading levels also appeared to be composed mostly of Ncap can be explained by incomplete denaturation. The contaminants identified were all from *E. coli* BL21 with the most frequent/abundant being Elongation factor Tu (EF-Tu, UNIPROT accession number A0A140NCI6). This is not surprising as EF-Tu is the most highly expressed gene product in *E. coli* [61]. The next highest-level contaminant was an uncharacterised protein (A0A140SS81), followed by a Type VI secretion system effector (A0A140N758), and an alcohol dehydrogenase GroES domain protein (A0A140N870), Transcriptional regulator LacI (A0A140NB96) which is encoded on the expression plasmid used in our work, methylmalonyl-CoA mutase (A0A140N835), sulphate ABC transporter (A0A140N7X7) and transcriptional regulator, IclR (A0A140NF03). All contaminants over 0.1% are shown in **Figure S6**.

### Response of Ncap variants with pooled antisera

We have previously validated an assay for the presence of human anti-SARS-CoV-2 Ncap antibodies using serum from SARS-CoV-2-confirmed cases and pre-pandemic serum samples [45]. We next compared the detectability of Ncap variants using pooled human sera (**Figure 5**).

**Figure 5.**
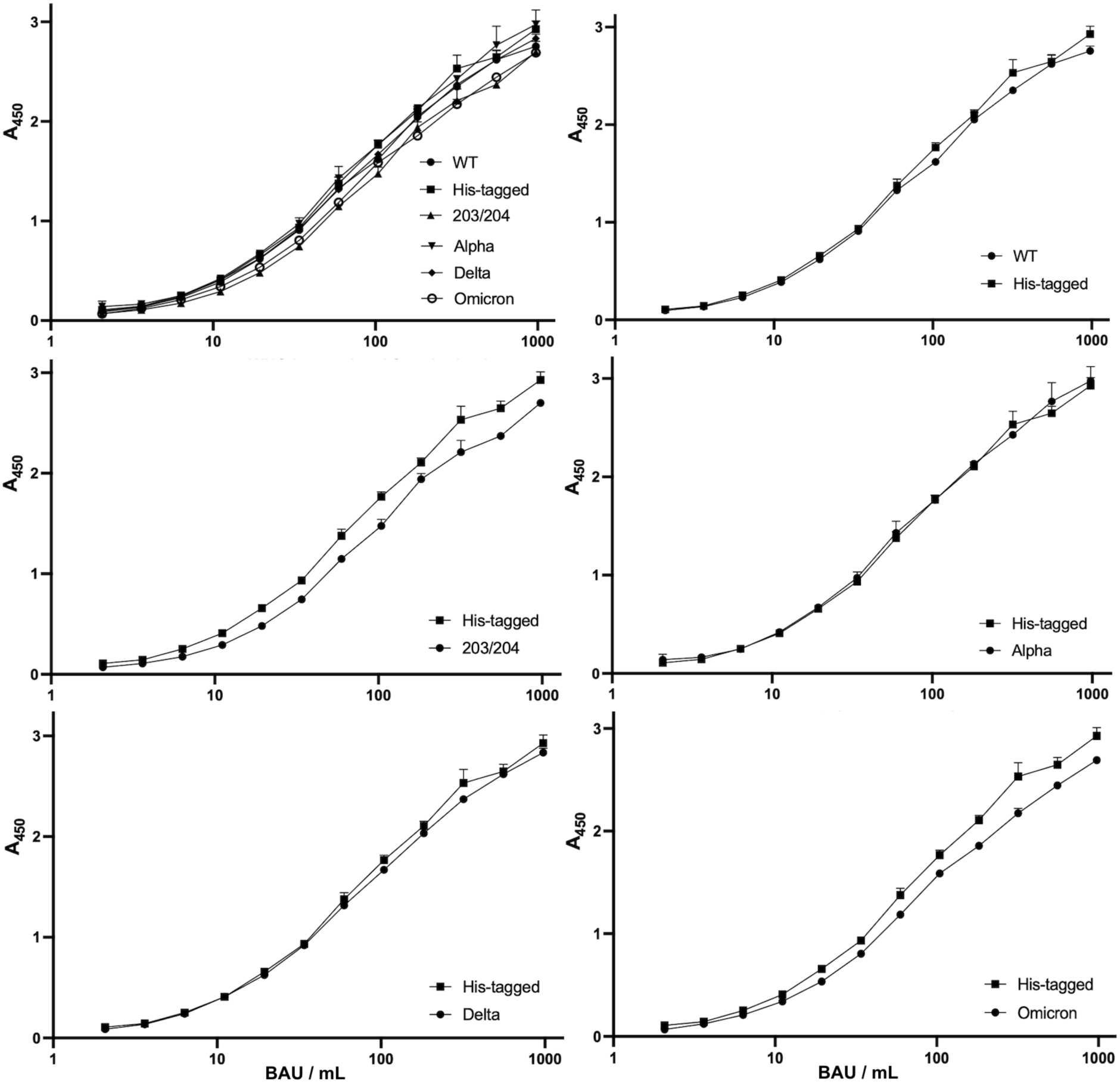
ELISA assay results for Ncap variants with pooled human sera. Ncap variants were immobilised on micro-titre plates and ELISA performed as described above (n=2). Top left panel compares all Ncaps produced (curve for B.1.1 is labelled 203/204). The original wild-type SARs-CoV-2 Ncap (untagged) and the his-tagged version are compared (top right). The remaining panels show pairwise comparisons between the latter and other 6His-tagged variants.

There was no significant difference between the measured ELISA signal response curves for the tagged and untagged original SARS-CoV-2 recombinant proteins, nor for the Delta or Alpha [62]. However, the Omicron and B.1.1 variants elicited lower ELISA signals from the pooled serum (P< 0.001 in both cases). For B.1.1 the median reduction was 18.5% (range 7.8–35.5%) while for Omicron, the median reduction was 14.0% (range 7.6–36.0%). The largest reduction in signal occurred at the lowest serum concentrations tested.

## CONCLUSION

We have developed generic protocols for production of soluble, immunologically competent Ncap proteins from *E. coli* without the need for denaturing conditions or refolding. The expression vector developed is portable as the synthetic T5 promoter is recognised by native sigma 70 of *E. coli* RNA polymerase [54] [63]. Thus, in principle it can be used in any *E. coli* strain unlike e.g. T7 expression systems which require the presence of the T7 RNA polymerase gene within the host genome or supplied on an additional genetic element [64]. Yields of up to 10 mg purified protein per gramme of cell paste were readily achieved, though we acknowledge that elements of the expression and purification protocols could be optimised for individual proteins to improve yields even further. All the molecules produced were recognised by pooled sera from confirmed SARS-CoV-2 convalescent patients although the B.1.1 and Omicron variants showed slightly reduced ELISA responses.

## Acknowledgements

We gratefully acknowledge the financial support of; Domen Zafred has received funding from the European Union’s Horizon 2020 research and innovation programme under the Marie Sklodowska-Curie grant agreement number 843245; a Wellcome Trust Intermediate Clinical Fellowship to T.d.S. (110058/Z/15/Z); BBSRC grants BB/V011456/1, BB/L015048/1 and EPSRC grants EP/T019328/1, EP/S01778X/1; Waters Corp. for ongoing support to the MBCCMS, EL is funded by a BBRSC DTP CASE award with Pharmaron Ltd. Infrastructure support funding for the Florey Institute AMR Research Capital Funding (grant number NIHR200636). We thank Zain Mahmood and Hiwa Khalis Jabar for supervised preliminary experiments on the Alpha and Delta variants and Dr Martin Nicklin for helpful discussions. Dr Adelina Acosta Martin kindly provided mass spectrometric analytical services (Faculty of Science biOMICS Facility with funding support from the Wolfson Foundation, Yorkshire Cancer Research, Leverhulme Trust, BBSRC, EPSRC, European Union, and the University of Sheffield Alumni fund). We acknowledge the support of the Mass Spectrometry and Separations Science facility at the University of Manchester and Reynard Spiess for help with instrumentation.

## Author Contributions

J.R.S., D.Z., P.E.B and T.d.S. acquired funding, designed and supervised the study and analysed data. E.L.B., M.B.P., D.Z., J.P., H.R.H., J. Darby, J. Dai, E.L., K.C. carried out experiments and analysed data. P.E.B, E.L.B., M.B.P., J.R.S. and D.Z. drafted the manuscript. All authors read and approved the final manuscript.

