## Supplementary Data, Figures and Tables for "Efficient overexpression and purification of SARS-CoV-2 Nucleocapsid proteins in *Escherichia coli*"

**Table of Contents**

| | | |
|---|---|---|
| **Table S1** | **Synthetic DNA used in this study** | **Page 2** |
| **Table S2** | **Synthetic oligonucleotides used in this study** | **Page 3** |
| **Table S3** | **Mass spectrometry analysis of purified Nucleocapsid proteins** | **Page 3** |
| **Table S4** | **Statistical analysis of Ncap ELISA responses** | **Page 3** |
| **Figure S1** | **Local impact of mutations on Ncap structure** | **Page 4** |
| **Figure S2** | **Molecular flexibility of Ncap and variants** | **Page 5** |
| **Figure S3** | **Mutations introduce changes in Ncap residue flexibility** | **Page 6** |
| **Figure S4** | **Purification of mini-His-tagged nucleocapsid proteins** | **Page 7** |
| **Figure S5** | **UV spectra of purified nucleocapsid proteins** | **Page 9** |
| **Figure S6** | **Contaminants in tagged and untagged nucleocapsid proteins** | **Page 10** |

### Table S1 Synthetic DNA used in this study

| Name /supplier | Sequence | Size / bps |
|---|---|---|
| **Ncap_N-terminal**<br>**GENEWIZ**<br>**Germany GmbH** | TATT**GAATTC**TAATTAATTAATAGAGGAGAAAAA**CATATG**CATCACCATCACCACCACGGATCAGATAAC<br>GGTCCACAAAACCAGCGTAACGCTCCGCGTATCACCTTCGGCGGCCCAAGCGATAGCACCGGTTCTAATC<br>AGAACGGTGAACGTAGCGGCGCGCGTTCCAAGCAACGTCGTCCGCAGGGCCTGCCGAACAACACGGCGTC<br>CTGGTTTACCGCTCTGACCCAGCATGGCAAAGA**AGATCT**GAAGTTCCCGCGTGGTCAAGGGGTGCCGATT<br>AACACCAACAGCTCGCCGGATGATCAAATCGGCTATTACCGCAGAGCAACCCGTCGTATCCGTGGCGGTG<br>ATGGTAAAATGAAAGACCTGAGCCCGCGTTGGTATTTCTACTACCTGGGTACAGGCCCGGAGGCTGGTCT<br>GCCGTATGGTGCCAATAAAGACGGTATTATCTGGGTCGCAACTGAAGGCGCGTTGAATACGCCGAAGGAC<br>CACATTGGTACTCGCAACCCGGCGAATAATGCGGCGATTGTACTGCAGTTGCCGCAGGGCACCACCCTGC<br>CGAAAGGTTTTTACGCAGAAGGTAGCCGTGGTGGCAGCCAGGCTTCGTCGCGCTCTAGTTCCCGCTCACG<br>CAACAGCAGCCGCAACAGCACCCCGGGTTCCTCCAGAGGTACGAGCCCGGCCCGTATGGCAGGCAACGGC<br>GGTGACGCTGCCCTTGCA | **718** |
| **Ncap_C-terminal**<br>**GENEWIZ**<br>**Germany GmbH** | CAACGGCGGTGACGCTGCCCTTGCACTGTTGTTGTTGGATCGTCTGAACCAACTTGAGAGCAAGATGTCC<br>GGTAAAGGCCAGCAACAGCAAGGTCAGACCGTGACCAAAAAGTCTGCGGCGGAGGCGAGTAAAAAACCGC<br>GACAGAAACGCACCGCGACCAAGGCGTATAACGTTACCCAAGCGTTTGGTCGTCGCGGTCCGGAGCAGAC<br>CCAGGGAAACTTCGGCGACCAAGAGCTGATTCGTCAGGGCACCGATTATAAGCATTGGCCGCAGATCGCC<br>CAGTTTGCACCATCAGCGTCTGCTTTCTTTGGCATGAGCCGTATCGGCATGGAAGTTACTCCTAGCGGCA<br>CTTGGCTGACGTACACCGGCGCCATCAAACTGGACGACAAGGACCCGAATTTCAAAGACCAAGTTATTCT<br>GCTGAATAAGCACATCGATGCGTACAAGACCTTCCCGCCTACGGAGCCGAAGAAGGACAAGAAAAAGAAG<br>GCAGATGAAACCCAAGCGTTGCCACAGCGTCAAAAAAAACAGCAAACGGTGACCCTGCTCCCGGCGGCAG<br>ACCTGGATGATTTTAGCAAGCAGTTACAACAAAGCATGTCTAGCGCTGACAGCACCCAAGCGTAATAA*CC*<br>*CGGGGCTAATAAGTCGCAAAAAACCCCGCTTCGGCGGGGTTTTTTCGCAAGC***AAGCTT**ATTT | **692** |
| **Alpha**<br>**NBS Biologicals**<br>**Ltd, UK** | **GAATTC**TAATTAATTAATAGAGGAGAAAACATATGCATCACCATCACCACCACGGATCACTGAACGGTCC<br>ACAAAACCAGCGTAACGCTCCGCGTATCACCTTCGGCGGCCCAAGCGATAGCACCGGTTCTAATCAGAAC<br>GGTGAACGTAGCGGCGCGCGTTCCAAGCAACGTCGTCCGCAGGGCCTGCCGAACAACACGGCGTCCTGGT<br>TTACCGCTCTGACCCAGCATGGCAAAGAAGATCTGAAGTTCCCGCGTGGTCAAGGGGTGCCGATTAACAC<br>CAACAGCTCGCCGGATGATCAAATCGGCTATTACCGCAGAGCAACCCGTCGTATCCGTGGCGGTGATGGT<br>AAAATGAAAGACCTGAGCCCGCGTTGGTATTTCTACTACCTGGGTACAGGCCCGGAGGCTGGTCTGCCGT<br>ATGGTGCCAATAAAGACGGTATTATCTGGGTCGCAACTGAAGGCGCGTTGAATACGCCGAAGGACCACAT<br>TGGTACTCGCAACCCGGCGAATAATGCGGCGATTGTACTGCAGTTGCCGCAGGGCACCACCCTGCCGAAA<br>GGTTTTTACGCAGAAGGTAGCCGTGGTGGCAGCCAGGCTTCGTCGCGCTCTAGTTCCCGCTCACGCAACA<br>GCAGCCGCAACAGCACCCCGGGTTCCTCCAAACGCACGAGCCCGGCCCGTATGGCAGGCAACGGCGGTGA<br>CGCTGCCCTTGCACTGTTGTTGTTGGATCGTCTGAACCAACTTGAGAGCAAGATGTTCGGTAAAGGCCAG<br>CAACAGCAAGGTCAGACCGTGACCAAAAAGTCTGCGGCGGAGGCGAGTAAAAAACCGCGACAGAAACGCA<br>CCGCGACCAAGGCGTATAACGTTACCCAAGCGTTTGGTCGTCGCGGTCCGGAGCAGACCCAGGGAAACTT<br>CGGCGACCAAGAGCTGATTCGTCAGGGCACCGATTATAAGCATTGGCCGCAGATCGCCCAGTTTGCACCA<br>TCAGCGTCTGCTTTCTTTGGCATGAGCCGTATCGGCATGGAAGTTACTCCTAGCGGCACTTGGCTGACGT<br>ACACCGGCGCCATCAAACTGGACGACAAGGACCCGAATTTCAAAGACCAAGTTATTCTGCTGAATAAGCA<br>CATCGATGCGTACAAGACCTTCCCGCCTACGGAGCCGAAGAAGGACAAGAAAAAGAAGGCAGATGAAACC<br>CAAGCGTTGCCACAGCGTCAAAAAAAACAGCAAACGGTGACCCTGCTCCCGGCGGCAGACCTGGATGATT<br>TTAGCAAGCAGTTACAACAAAGCATGTCTAGCGCTGACAGCACCCAAGCGTAAT**AAGCTT** | **1320** |
| **Delta**<br>**NBS Biologicals**<br>**Ltd, UK** | **GAATTC**TAATTAATTAATAGAGGAGAAAACATATGCATCACCATCACCACCACGGATCAGATAACGGTCC<br>ACAAAACCAGCGTAACGCTCCGCGTATCACCTTCGGCGGCCCAAGCGATAGCACCGGTTCTAATCAGAAC<br>GGTGAACGTAGCGGCGCGCGTTCCAAGCAACGTCGTCCGCAGGGCCTGCCGAACAACACGGCGTCCTGGT<br>TTACCGCTCTGACCCAGCATGGCAAAGAAGGTCTGAAGTTCCCGCGTGGTCAAGGGGTGCCGATTAACAC<br>CAACAGCTCGCCGGATGATCAAATCGGCTATTACCGCAGAGCAACCCGTCGTATCCGTGGCGGTGATGGT<br>AAAATGAAAGACCTGAGCCCGCGTTGGTATTTCTACTACCTGGGTACAGGCCCGGAGGCTGGTCTGCCGT<br>ATGGTGCCAATAAAGACGGTATTATCTGGGTCGCAACTGAAGGCGCGTTGAATACGCCGAAGGACCACAT<br>TGGTACTCGCAACCCGGCGAATAATGCGGCGATTGTACTGCAGTTGCCGCAGGGCACCACCCTGCCGAAA<br>GGTTTTTACGCAGAAGGTAGCCGTGGTGGCAGCCAGGCTTCGTCGCGCTCTAGTTCCCGCTCACGCAACA<br>GCAGCCGCAACAGCACCCCGGGTTCCTCCATGGGTACGAGCCCGGCCCGTATGGCAGGCAACGGCTGTGA<br>CGCTGCCCTTGCACTGTTGTTGTTGGATCGTCTGAACCAACTTGAGAGCAAGATGTCCGGTAAAGGCCAG<br>CAACAGCAAGGTCAGACCGTGACCAAAAAGTCTGCGGCGGAGGCGAGTAAAAAACCGCGACAGAAACGCA<br>CCGCGACCAAGGCGTATAACGTTACCCAAGCGTTTGGTCGTCGCGGTCCGGAGCAGACCCAGGGAAACTT<br>CGGCGACCAAGAGCTGATTCGTCAGGGCACCGATTATAAGCATTGGCCGCAGATCGCCCAGTTTGCACCA<br>TCAGCGTCTGCTTTCTTTGGCATGAGCCGTATCGGCATGGAAGTTACTCCTAGCGGCACTTGGCTGACGT<br>ACACCGGCGCCATCAAACTGGACGACAAGGACCCGAATTTCAAAGACCAAGTTATTCTGCTGAATAAGCA<br>CATCGATGCGTACAAGACCTTCCCGCCTACGGAGCCGAAGAAGGACAAGAAAAAGAAGGCATATGAAACC<br>CAAGCGTTGCCACAGCGTCAAAAAAAACAGCAAACGGTGACCCTGCTCCCGGCGGCAGACCTGGATGATT<br>TTAGCAAGCAGTTACAACAAAGCATGTCTAGCGCTGACAGCACCCAAGCGTAAT**AAGCTT** | **1320** |
| **Omicron**<br>**GENEWIZ**<br>**Germany GmbH** | TTTA**GAATTC**TAATTAATTAATAGAGGAGAAAAACATATGCATCACCATCACCACCACGGATCAGATAAC<br>GGTCCACAAAACCAGCGTAACGCTCTGCGTATCACCTTCGGCGGCCCAAGCGATAGCACCGGTTCTAATC<br>AGAACGGTGGCGCGCGTTCCAAGCAACGTCGTCCGCAGGGCCTGCCGAACAACACGGCGTCCTGGTTTAC<br>CGCTCTGACCCAGCATGGCAAAGA**AGATCT**GAAG | **244** |

**Table S1 Footnote** Recognition sites for restriction endonucleases used for cloning (EcoRI, GAATTC; NdeI, CATATG; BglII, AGACTC and HindIII, AAGCTT) are underlined in bold. The 25 bp overlap between N-terminal and C-terminal coding regions of SARS-CoV-2 Ncap fragments are highlighted in cyan.

**Table S2 Synthetic oligonucleotides used in this study**

| | | |
|---|---|---|
| **Ncap_for** | 5' - AAATGAATTCTAAATAATTAATAGAGGAGAAAAAC - 3' | **35** |
| **Ncap_rev** | 5' - AATAAAGCTTATTACGCTTGGGTGCTGTCAGC - 3' | **32** |
| **Native_Ncap_for** | 5' - GAAAAACATATGTCAGATAACGGTCCACAAAACC - 3' | **34** |
| **Ncap_203/204_for** | 5' - GGTTCCTCC**AAACGC**ACGAGCCCGGCCCGT - 3' | **30** |
| **Ncap_203/204_rev** | 5' - GCTCGT**GCGTTT**GGAGGAACCCGGGGTGCTGTT - 3' | **33** |

Recognition sites for restriction endonucleases used for cloning (EcoRI, GAATTC; NdeI, CATATG and HindIII, AAGCTT) are highlighted in black. Nucleotides introducing the 203/204 Lys/Gly to Arg/Lys are marked in bold text.

#### Table S3 Mass spectrometry analysis of purified Nucleocapsid proteins

| Protein | Mass Expected | Mass Observed |
|---|---|---|
| Ncap (WT) | 45495.5 | 45495.8 |
| MHT Ncap (WT) | 46505.6 | 46507.3 |
| MHT Ncap (B.1.1) | 46576.7 | 46577.0 |
| MHT Alpha | 46634.9 | 46633.0 |
| MHT Delta | 46516.7 | 46516.5 |
| MHT Omicron | 46220.4 | 46220.2 |

#### Table S4 Statistical analysis of Ncap ELISA responses

| | Ncap Untagged | MHT Ncap | MHT B.1.1 | MHT Alpha | MHT Delta | MHT Omicron |
|---|---|---|---|---|---|---|
| **Ncap Untagged** | | | | | | |
| **MHT Ncap** | 0.105 | | | | | |
| **MHT B.1.1** | $6.21 \times 10^{-7}$ | $1.40 \times 10^{-9}$ | | | | |
| **MHT Alpha** | $8.82 \times 10^{-3}$ | 0.954 | $1.36 \times 10^{-7}$ | | | |
| **MHT Delta** | 1.00 | 0.031 | $8.07 \times 10^{-10}$ | $6.39 \times 10^{-2}$ | | |
| **MHT Omicron** | $8.52 \times 10^{-5}$ | $2.01 \times 10^{-7}$ | $1.30 \times 10^{-3}$ | $1.06 \times 10^{-6}$ | $2.28 \times 10^{-8}$ | |

Uncorrected P values shown [1]. Greyed values remain statistically significant after applying a strict Bonferroni correction for multiple testing [2].

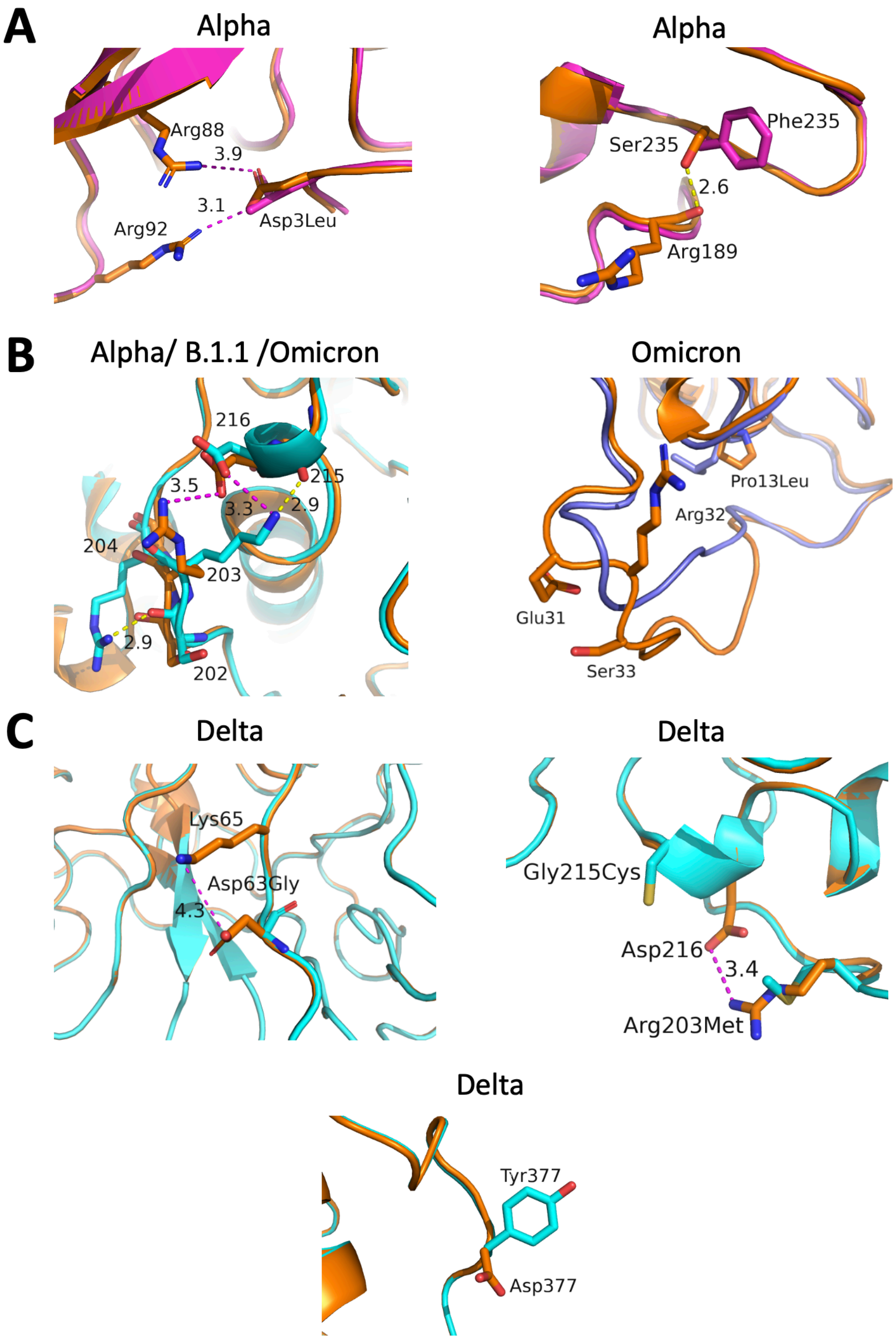


**Figure S1 Local impact of mutations on Ncap structure**
Wild-type Ncap in gold cartoon backbones with relevant side chains in sticks are shown compared with variant proteins. Charge-charge (ion pair) interactions are shown as magenta dotted lines and hydrogen bonds indicated by dotted yellow lines. Distances are indicated in Å. **(A)** The Asp3Leu mutation in Alpha (magenta cartoon) results in the loss of two ion-pair interactions (magenta dotted lines) with the residues shown. The Ser235Phe substitution results in a loss of a hydrogen bond with the backbone carbonyl of Arg189. **(B)** The Arg203Lys–Gly204Arg double substitution present in B.1.1, Alpha and Omicron maintains an ion pair interaction with Aps216 and introduces a possible hydrogen bond with the backbone of the same residue. The Omicron Pro13Leu and 31-33 deleted residues are depicted in the right-hand panel. **(C)** Modelling of substitutions in Delta suggest loss of two ion-pairs (Lys65–Asp63 at Asp63Gly and Asp216–Arg203 at Arg203Met).

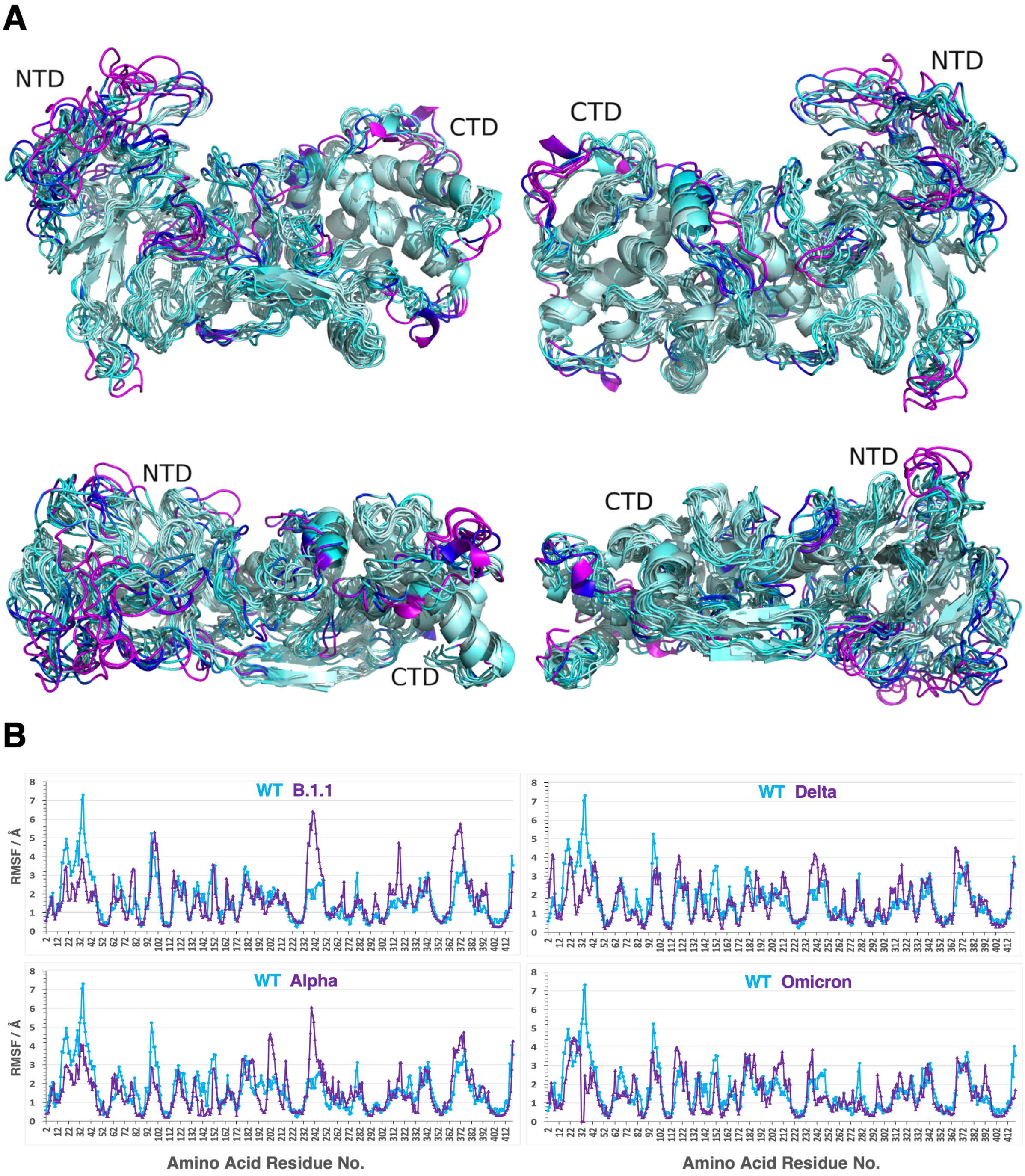


**Figure S2 Molecular flexibility of Ncap and variants**

**(A)** Views of ensemble of structures of Ncap obtained using molecular dynamics simulation rendered in PyMOL, coloured by root mean square flexibility (RMSF)generated by CABSFLEX molecular dynamics server [3]. The most rigid residues are coloured pale cyan followed by aquamarine then cyan then blue with magenta being the most flexible. **(B)** The predicated changes in RMSF for each residue in individual NCAP variants (magenta) is compared with the WT protein (cyan).

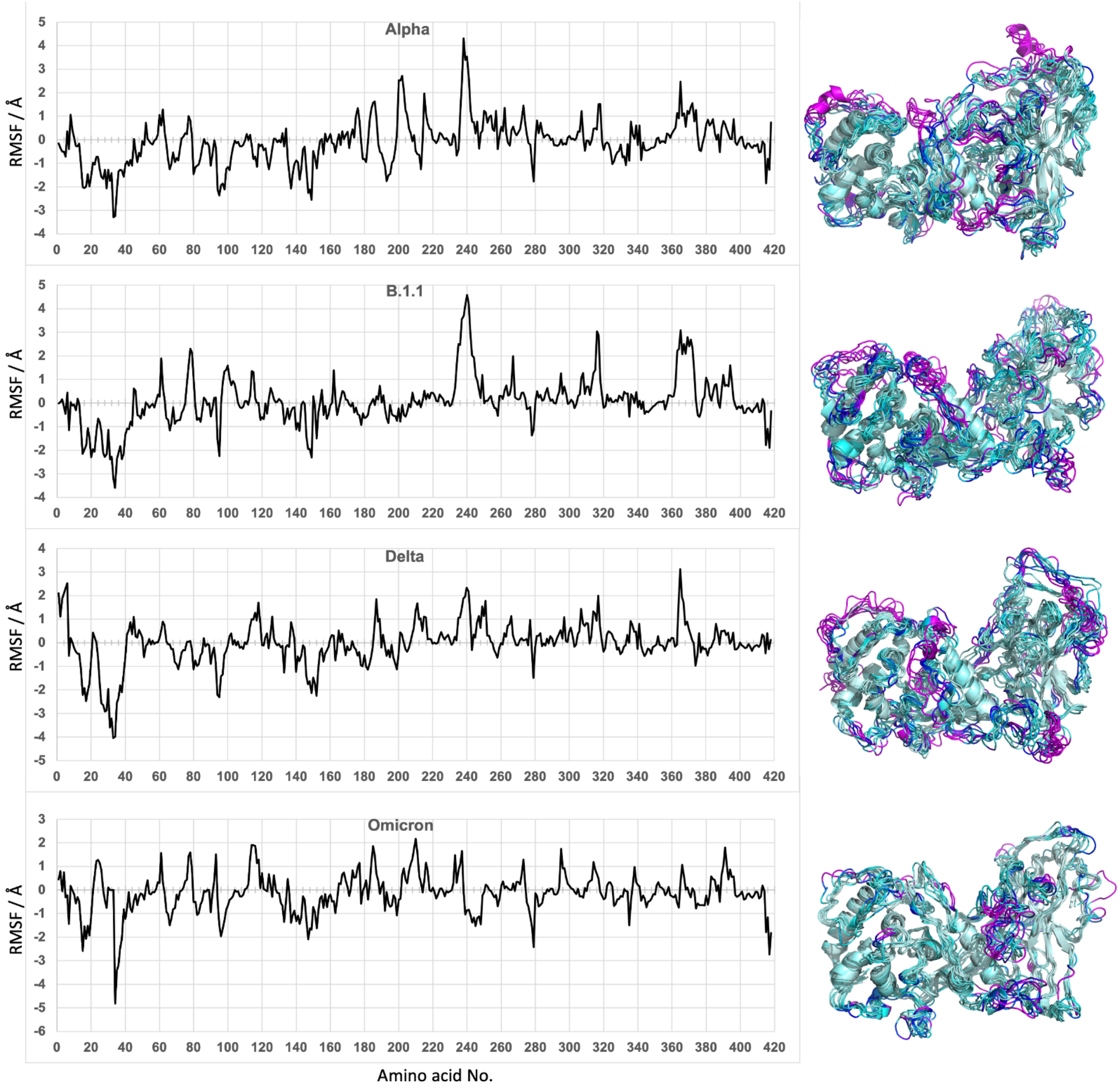


**Figure S3 Mutations introduce changes in Ncap residue flexibility**
The mean flexibility changes observed in variants compared to the original Ncap calculated from the structures shown in **Figure S2** are shown on the left. Increased flexibility compared to wild type is indicated by positive values. Molecular graphics shown on the right for each variant present MD trajectories coloured by RMSF as in **Figure S2**.

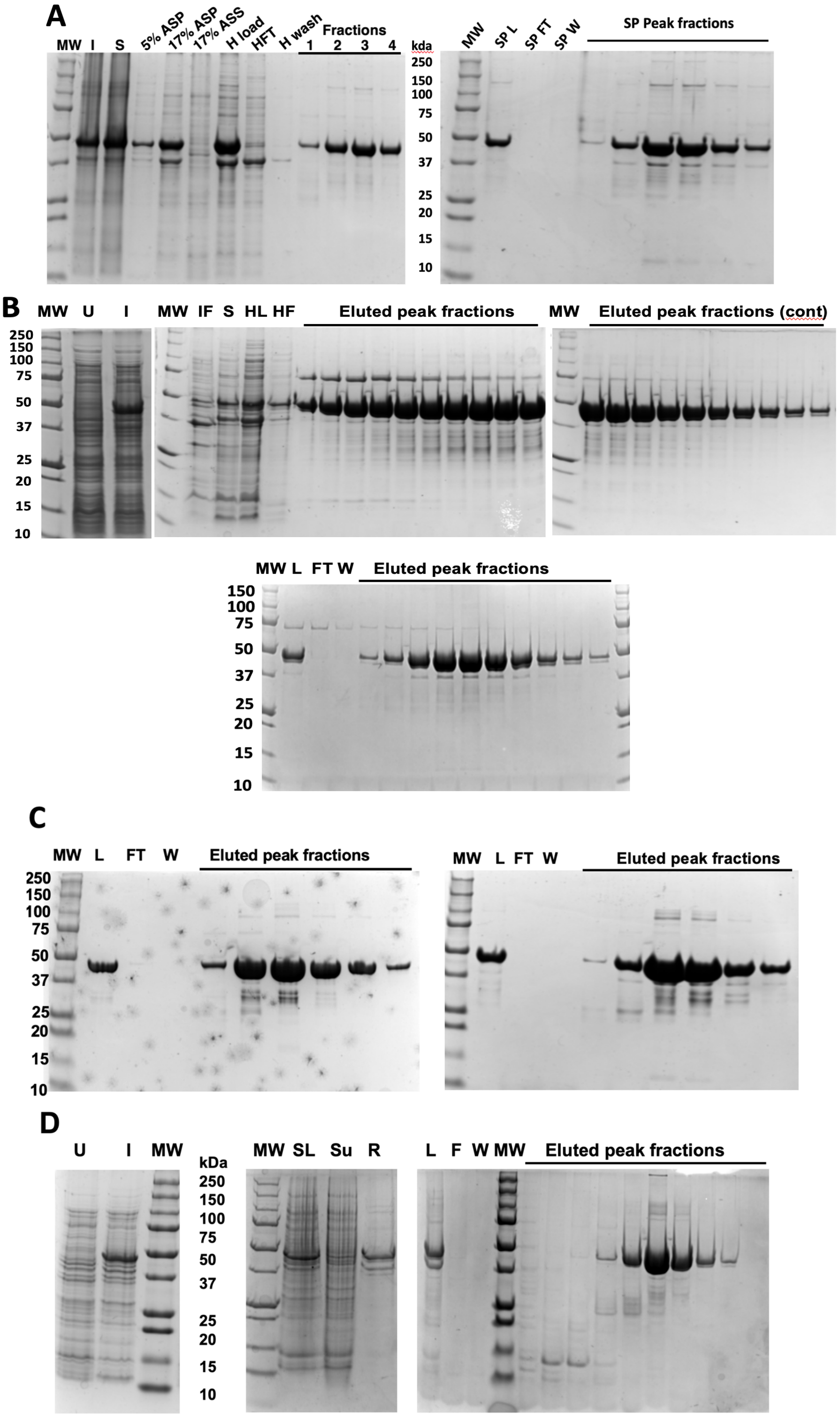


### Figure S4 Purification of Ncap proteins

**Panel A** Purification of mini-His-tagged Delta Ncap monitored by Coomassie stained SDS-PAGE. BioRad standard molecular weight markers are shown on both gels (MW), sizes indicated in centre. Left panel shows expression (Lane I) induced, (Lane S) soluble cell lysates. Insoluble material precipitated by 5% and 17% ammonium sulphate (Lane 5 & 17% ASP). Material remaining soluble in 17% ammonium sulphate (Lane 17% ASS). Lanes H Load, HFT and H wash show protein applied to HisTrap nickel chelate column and peak eluted fractions number 1- 4. Pooled fractions from the chelate column, loaded onto an SP-Sepharose cation exchange column after 10-fold dilution (Lane SP L, Right panel). The flow through, wash and eluted proteins from the salt gradient are shown in Lanes SP FT, SP wash, and SP peak fractions, respectively. **Panel B** Outlines purification of B.1.1 MHT-Ncap; uninduced (U), induced (I) , insoluble (IF), soluble (S) and load onto HisTrap column (HL) flow through (HF), peak fractions shown in the upper three gels. Lower central image shows load (L), flow through (FT), wash (W) and fractions eluted from SP Sepharose column. **Panel C** Shows latter stages of purification of Alpha MHT-Ncap. Load (L), flow through (FT), wash (W) and fractions eluted from the HisTrap column on left gel. Load (L), flow through (FT), wash (W) and fractions eluted from the SP Sepharose column on the right-hand side. **Panel D** Shows whole-cell lysate of uninduced and induced cells expressing untagged Ncap (U and I, in left image), soluble lysate from induced cells (SL), soluble supernatant and resuspended pellet (Su and R respectively) from 17% ammonium sulphate precipitation. The latter was passed through a Q FF anion exchanger and the flow through (L, right hand image) loaded on to an SP anion exchange column. Samples of flow through (F), wash (W) and eluted peak samples are shown.

**Figure S5 UV spectra**

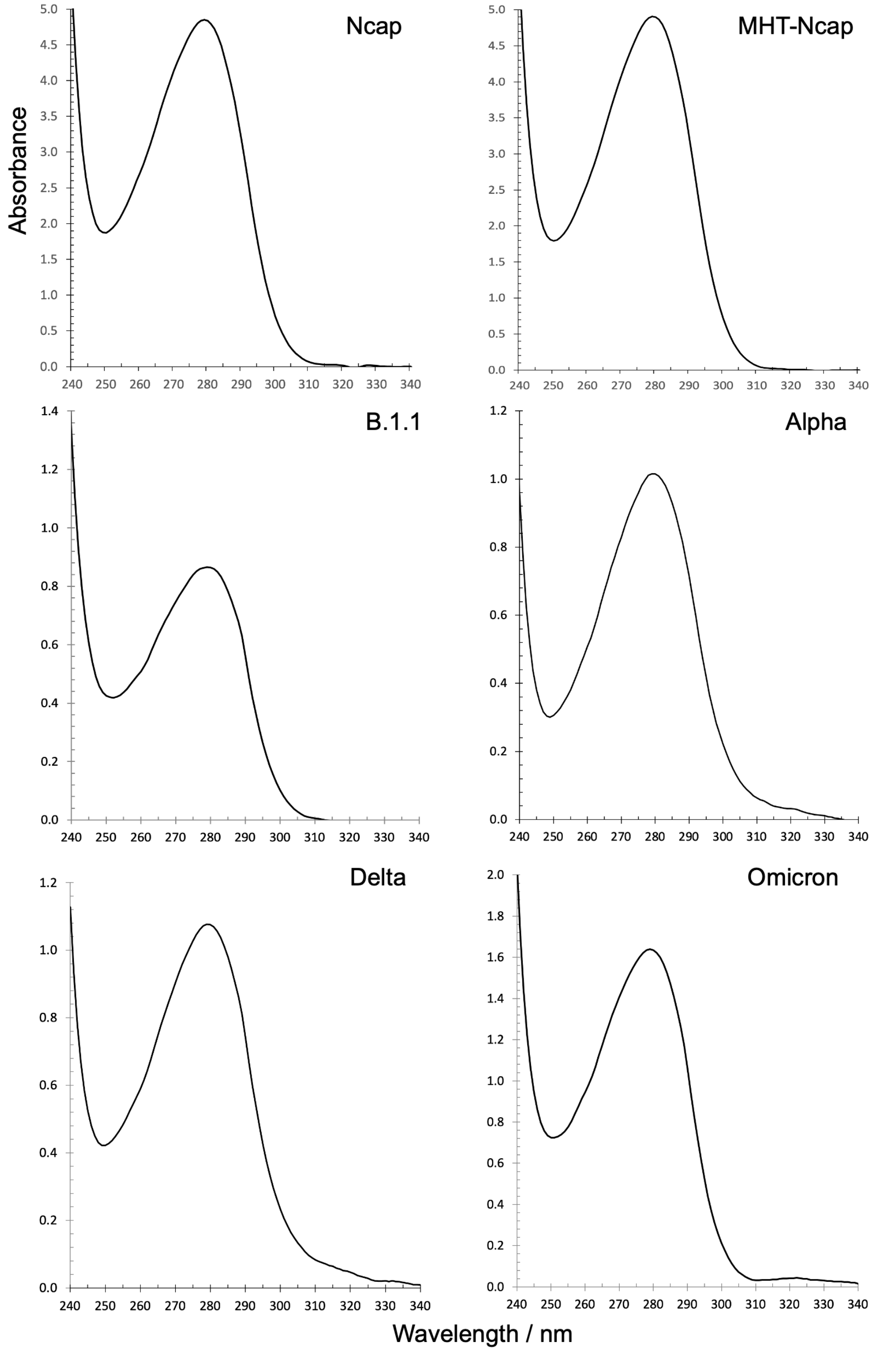


**Figure S5 UV spectra of purified Nucleocapsid proteins**
UV spectra of purified proteins were recorded using an IMPLEN NanoPhotometer N60. The 260 / 280 nm ratios calculated varied from 0.500 – 0.59 indicating little contamination with nucleic acid.

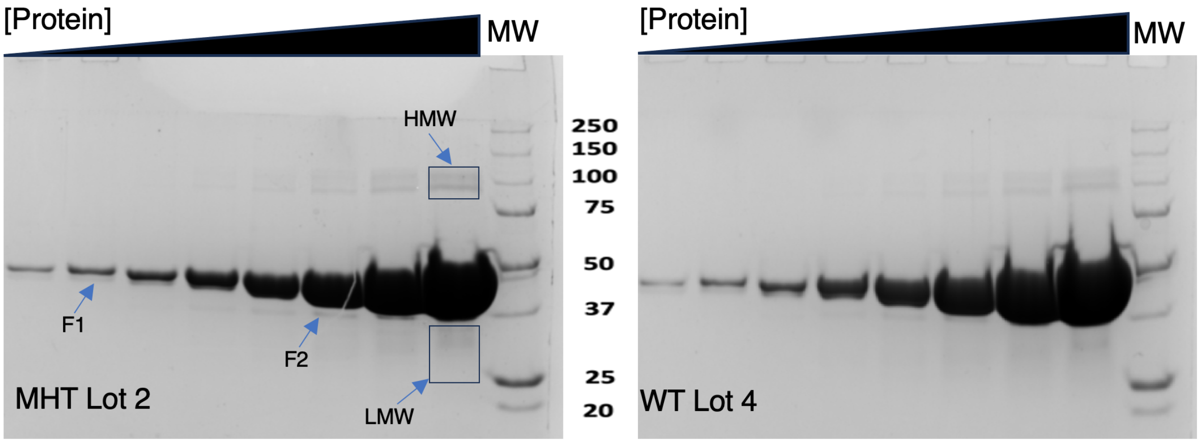


| Main Band/Fragment 1 | % | Main Band/Frgament 1 | % |
|---|---|---|---|
| MHT-NCP | 97.67 | WT-NCP | 98.03 |
| A0A140SS81 Uncharacterized protein | 1.62 | A0A140N835 Methylmalonyl-CoA mutase | 1.06 |
| A0A140NCI6 Elongation factor Tu | 0.10 | C5W7S9 tRNA1 -methyltransferase | 0.49 |
| | | A0A140NCM1 O-succinylhomoserine (Thiol)-lyase | 0.37 |
| **Bands in HMW group** | | **Bands in HMW group** | |
| MHT-NCAP | 96.46 | WT-NCP | 97.76 |
| A0A140NCI6 Elongation factor Tu | 0.49 | A0A140N7X7 Sulfate ABC transporter | 0.64 |
| A0A140N870 Alcohol dehydrogenase GroES domain | 0.40 | A0A140NB96 Transcriptional regulator, LacI | 0.41 |
| A0A140N758 Type VI secretion system effector | 0.28 | A0A140N758 Type VI secretion system effector, Hcp1 | 0.27 |
| A0A140NB96 Transcriptional regulator, LacI | 0.25 | A0A140NF68 Cys-tRNA(Pro/Cys) deacylase | 0.10 |
| A0A140NH65 Chaperonin GroEL | 0.24 | | |
| A0A140NFV3 Chaperone protein DnaK | 0.17 | | |
| A0A140N7K8 Small ribosomal subunit protein uS14 | 0.11 | | |
| A0A140N403 YceK/YidQ family lipoprotein | 0.11 | | |
| A0A140N951 NADH-ubiquinone oxidoreductase chain 4L | 0.11 | | |
| A0A140NC97 Alkyl hydroperoxide reductase C | 0.10 | | |
| **Fragment 2** | | **Fragment 2** | |
| MHT-NCP | 91.46 | WT-NCP | 96.62 |
| A0A140SS81 Uncharacterized protein | 6.46 | A0A140N7X7 Sulfate ABC transporter | 1.61 |
| A0A140N758 Type VI secretion system effector | 0.59 | A0A140N638 Fructose-1,6-bisphosphatase | 0.42 |
| A0A140N870 Alcohol dehydrogenase GroES domain | 0.32 | A0A140N582 Cell shape-determining protein MreB | 0.28 |
| A0A140NC78 PTS system, N-acetylglucosamine-specific | 0.31 | A0A140NB96 Transcriptional regulator, LacI family | 0.24 |
| A0A140NB96 Transcriptional regulator, LacI family | 0.25 | A0A140N960 Uncharacterized protein | 0.17 |
| A0A140NHP0 Branched-chain-amino-acid aminotransferase | 0.21 | | |
| **LMW smear** | | **LMW Smear** | |
| MHT-NCP | 96.16 | WT-NCP | 93.73 |
| A0A140N758 Type VI secretion system effector | 1.25 | A0A140NF03 Transcriptional regulator, IclR | 2.29 |
| A0A140NEJ5 3,4-dihydroxyphenylacetate 2,3-dioxygenase | 0.35 | A0A140N7X7 Sulfate ABC transporter | 1.81 |
| A0A140N8Y5 Glutaredoxin | 0.35 | A0A140NAZ8 Polyamine aminopropyltransferase | 0.44 |
| A0A140NAE2 Alpha-2-macroglobulin domain protein | 0.30 | A0A140NB54 RNA chaperone ProQ | 0.38 |
| A0A140SSC0 Thioredoxin | 0.26 | A0A140NC97 Alkyl hydroperoxide reductase C | 0.22 |
| A0A140N870 Alcohol dehydrogenase GroES domain | 0.20 | A0A140NCL7 Pseudouridine synthase | 0.15 |
| A0A140NB96 Transcriptional regulator, LacI family | 0.16 | A0A140NC78 PTS system | 0.15 |
| A0A140NCI6 Elongation factor Tu | 0.13 | A0A140NCM1 O-succinylhomoserine (Thiol)-lyase | 0.11 |
| | | A0A140N8L0 FAD-dependent pyridine nucleotide-disulphide oxidoreductase | 0.11 |

**Figure S6 Contaminants in tagged and untagged Ncap proteins**
Upper left panel shows increasing amounts of mini-His-tagged Ncap (MHT Lot2) loaded on a 10% SDS-PAGE gel allowing visualisation of contaminants. Lanes 1 through 8 are loaded with 0.5, 1, 2, 5, 10, 25, 50 and 100 µg of purified protein, respectively. Lane MW shows size markers with molecular weights in kDa indicated (BioRad, #161-0362). Contaminant bands are indicated by the arrows and boxes and include two higher MW (HMW) bands, as well as bands F1 and F2 running slightly ahead of the main protein band and a faint smear of low MW (LMW) bands running between the 37 and 25 kDa MW markers. Similar sized bands are seen in the Ncap (untagged, wild-type, WT Lot 4), upper right panel. Lower panel shows identities of the bands observed for contaminants over 0.1% according to quantification by iBAQ [4]. Process contaminants (mostly keratins) were also observed but were excluded from the analysis.
